# Mitochondrial mass governs the extent of T cell senescence

**DOI:** 10.1101/627240

**Authors:** Lauren A. Callender, Elizabeth C. Carroll, Emilia A Bober, Arne N. Akbar, Egle Solito, Sian M. Henson

## Abstract

The susceptibility of human CD4^+^ and CD8^+^ T cells to senesce differs, with CD8^+^ T cells acquiring an immunosenescent phenotype faster than their CD4^+^ T cell compartment. We show here that it is the inherent difference in mitochondrial content that drives this phenotype, with senescent human CD4^+^ T cells displaying a higher mitochondrial mass. The loss of mitochondria in the senescent human CD8^+^ T cells has knock-on consequences for nutrient usage, metabolism and function. Mitochondrial dysfunction has been linked to both cellular senescence and ageing, however it is still unclear whether mitochondria play a causal role in senescence. Our data shows that reducing mitochondrial function in human CD4^+^ T cells, through the addition of low dose rotenone, causes the generation of a CD4^+^ T cell with a CD8^+^ -like phenotype. Therefore we wish to propose that it is the inherent metabolic stability that governs the susceptibility to an immunosenescent phenotype.

## Introduction

The human immune system functionality declines with age in a process referred to as immunosenescence. The functional outcomes of this process include the compromised ability of older individuals to mount protective immune responses against both previously encountered and new pathogens^1^. Additionally, there is a marked decrease in vaccine efficacy in these populations^1^. While these age-associated alterations arise from defects in different leukocyte populations, the dysfunction is most profound in T cell subsets^2^. Furthermore, ageing is associated with a chronic low grade inflammatory state, termed inflammaging^3^ and mediates an important role in a range of age-related degenerative pathologies^4^. The source of this inflammation has yet to be defined. Senescent T cells are found to accumulate with age and represent a likely contributor to this inflammatory state that is observed during ageing^1^.

Primary human senescent T cells are a highly differentiated subset of cells found within the CD27^−^ CD28^−^ population^5, 6^. This subset can be further characterised on the basis of CD45RA expression, with highly differentiated T cells that re-express CD45RA identified as the senescent T cell population (EMRA; effector memory CD45RA re-expressing T cells). They display multiple characteristics of senescence including a low proliferative activity, high levels of DNA damage and loss of telomerase activity^7, 8^. However the response patterns of CD4^+^ and CD8^+^ T cells to ageing differs, with CD8^+^ T cells being more susceptible to both phenotypic and functional changes during ageing^9, 10^. The CD8^+^ EMRA T cell subset accumulate in higher proportions with age and are more prevalent following in vitro culture than the CD4^+^ EMRAs^9^. The cause of this difference has been suggested to be due to the differing homeostatic mechanisms and an increased gene expression instability of regulatory cell surface molecules in the CD8^+^ EMRA subset^9^. We would like to postulate an alternate view, that CD4^+^ EMRA T cells are more metabolically robust and therefore better able to withstand the intrinsic or extrinsic effects governing differentiation.

Metabolic examination of CD4^+^ and CD8^+^ T cells suggest that their metabolic programming allows differential immunological functions to be performed. We demonstrate here that CD4^+^ T cells have a greater mitochondrial mass and are consistently more oxidative than CD8^+^ T cells, allowing them to sustain effector function. Whereas the metabolic programs that prioritise rapid biosynthesis such as glycolysis are favoured by CD8^+^ T cells, allowing for faster growth and proliferative rates^11, 12^. We have previously shown that CD8^+^ EMRA T cells display impaired mitochondrial function^8^ but are still unclear as to whether CD4^+^ EMRA T cells also exhibit mitochondrial dysfunction. We provide evidence that this is not the case, CD4^+^ EMRA T cells have fitter, healthier mitochondria that are better able to meet the energy requirements of the CD4^+^ EMRA subset. Therefore we propose that it is the inherent metabolic stability that governs the susceptibility to an immunosenescent phenotype.

## Results

### Human CD4^+^ EMRA T cells development at a slower rate due to their higher mitochondrial content

Human T cells can be subdivided into four populations on the basis of their relative surface expression of CD45RA and CD27 molecules (Supplementary Figure 1A). The four subsets are defined as, naïve (N; CD45RA^+^CD27^+^), central memory (CM; CD45RA^−^CD27^+^), effector memory (EM; CD45RA^−^CD27^−^), and effector memory T cells that re-express CD45RA (EMRA; CD45RA^+^CD27^−^). We and others have demonstrated that the EMRA population exhibit numerous characteristics of senescence^8, 13, 14^, indeed it has also been known for some time that CD4^+^ EMRA T cells senesce at a slow rate than their CD8^+^ counterparts (Figure 1A). It was thought that the difference in EMRA accumulation was due to their differing cytokine stabilities^9^, however we now demonstrate that it is a difference in mitochondrial mass between CD4^+^ and CD8^+^ EMRAs that governs the rate at which they develop.

**Figure 1.**
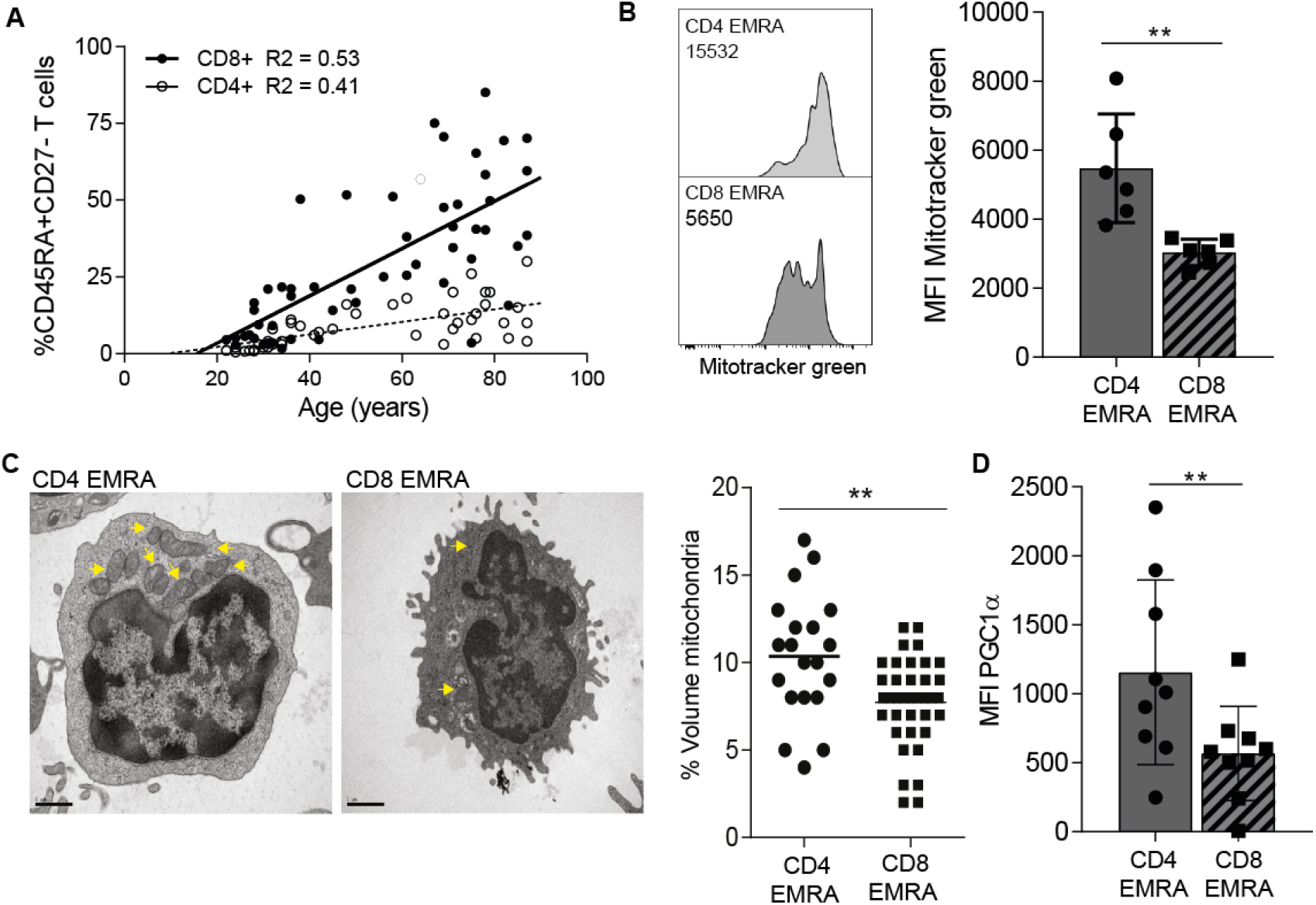
Human CD4^+^ EMRA T cells are acquired at a slower rate owing to a higher degree of mitochondrial content. (A) The accumulation of senescent CD4^+^ and CD8^+^ T cells with age defined by the markers CD45RA and CD27. (B) Representative flow cytometry plots and cumulative graphs of mitotracker green staining in CD4^+^ and CD8^+^ EMRA T cells analysed directly *ex vivo*. Data expressed as mean ± SEM of 6 donors. (C) Electron microscope images of CD4^+^ and CD8^+^ EMRA T cells imaged directly *ex vivo*. Yellow arrows mark mitochondria. Graph shows the percentage by cell volume of mitochondria in senescent T cell subsets determined by a point counting grid method from 20 different electron microscope images. (D) PGC1α expression in CD45RA/CD27 defined EMRA T cell subsets. Data expressed as mean ± SEM of 9 donors. p-values were calculated using a T-test.

Using MitoTracker Green, a mitochondrial-specific dye that binds the mitochondrial membranes independently of mitochondrial membrane potential (MMP), we found the CD4^+^ EMRA subset to have a significantly higher mitochondrial mass than CD8^+^ EMRAs, nearly double the amount of mitochondrial content (Figure 1B). The CD4^+^ EMRA subset retain their mitochondrial content copared to earlier less differentiated subsets (Supplementary Figure 2A) where as the CD8^+^ EMRAs do not^8^. This was also borne out when the EMRA subsets were examined ex-vivo by electron microscopy. We observed significantly fewer mitochondrial in the CD8^+^ EMRA compartment when compared to the CD4^+^ EMRA fraction using a point counting method (Figure 1C). Furthermore when we investigated the expression of PGC1α (peroxisome proliferator-activated receptor gamma coactivator 1-alpha), the key regulator of mitochondrial biogenesis, the CD4^+^ EMRA subset showed significantly higher ex-vivo levels of this marker than the CD8^+^ EMRAs (Figure 1D). Collectively our results demonstrate that senescent CD4^+^ T cells have increased mitochondrial mass in comparison to their CD8^+^ counterparts.

### Distinct mitochondrial functions in CD4^+^ and CD8^+^ EMRA subsets

The increased mitochondrial mass seen in the CD4^+^ EMRA subsets suggest they may exhibit distinct mitochondrial functions compared to the CD8^+^ EMRAs. Indeed using TMRE, which measures mitochondrial transmembrane potential, we found the CD4^+^ EMRAs had a higher proportion of hyperpolarised mitochondria than the CD8^+^ EMRA subset, which displayed a hypopolarised phenotype (Figure 2A). Hyperpolarised mitochondria can be a source of reactive oxygen species (ROS), which has the potential to drive senescence^15^. Mitochondrial ROS, measured using MitoSox, was found to be significantly higher in CD4^+^ EMRAs (Figure 2B), however this increase was neutralised owing to the higher mitochondrial mass, meaning that CD8^+^ EMRA T cells produced more ROS per mitochondria that can potentially enhance their senescent phenotype (Figure 2C). Furthermore increased ROS can also cause DNA damage and the activation of the DNA damage response, elevated during senescence. The examination of phosphorylated H2AX (γH2AX), a member of the histone H2A family that is part of the DNA damage response, in EMRAs revealed that the CD8^+^ EMRA subset displayed a higher level of this marker compared to CD4^+^ EMRAs (Figure 2D). However both EMRA subsets express the highest amount of DNA damge compared to their less differentiaited subsets (data not shown). We suggest that a loss of mitochondrial mass in the CD8^+^ EMRA subsets is a key mediator in generating an enhanced senescent state.

**Figure 2.**
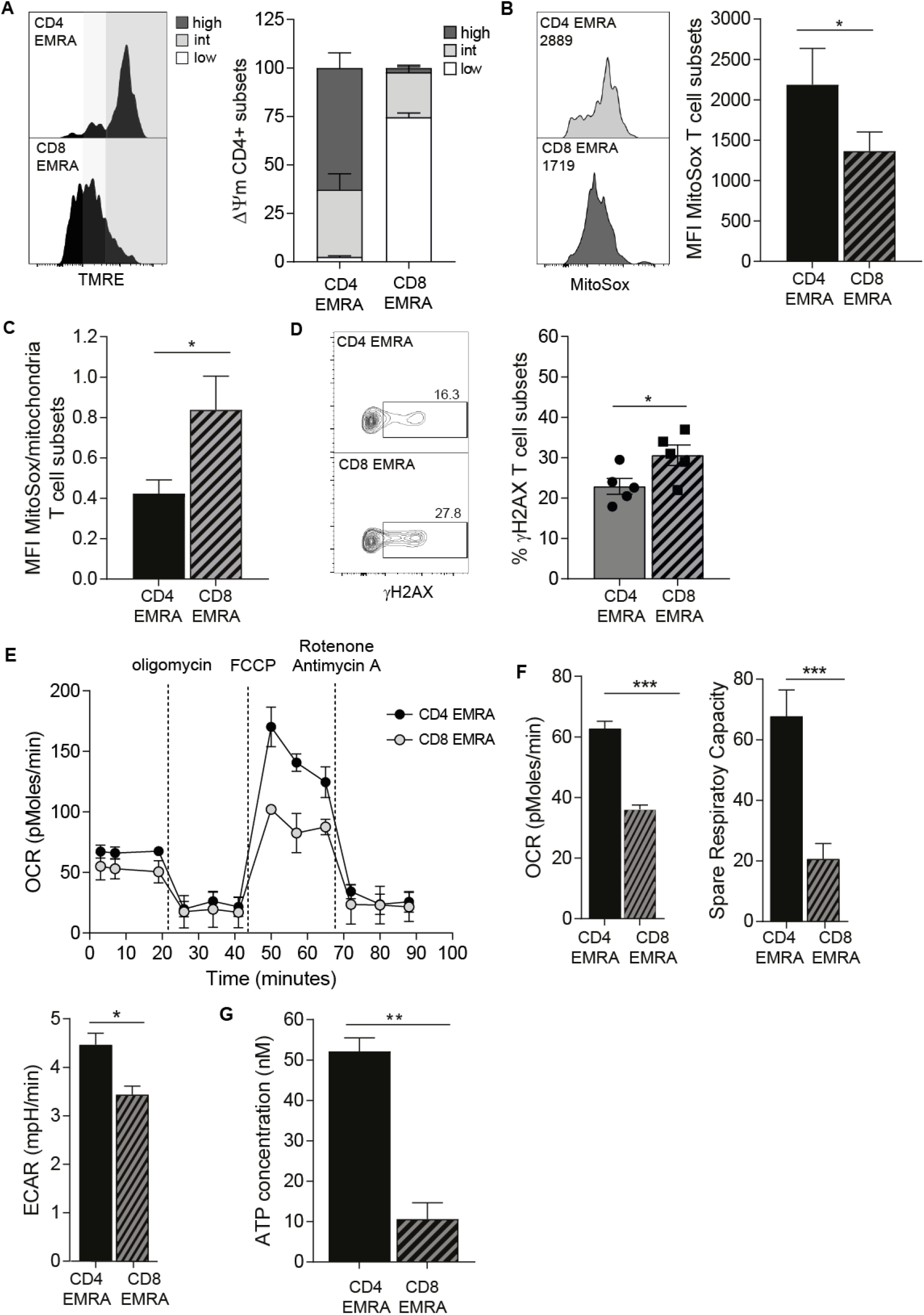
Mitochondrial dysfunction is observed CD8^+^ but not CD4^+^ EMRA T cell subsets. (A) Representative flow cytometry plots and cumulative graphs of TMRE staining showing membrane potential in CD45RA/CD27 T cell subsets directly *ex vivo*. defined showing the percentage of cortactin positive (a) CD4^+^ and (b) CD8^+^ T cells analysed directly *ex vivo*. Data expressed as mean ± SEM of 6 donors. (B) Mitochondrial ROS measured using MitoSox by flow cytometry in CD4^+^ and CD8^+^ EMRA T cells. Data expressed as mean ± SEM of 6 donors. (C) Mitochondrial ROS production expressed as a ratio of mitochondrial mass. (D) γH2AX expression as determined by flow cytometry in CD45RA/CD27 defined T cell subsets directly *ex vivo*, the graph shows the mean ± SEM for 5 donors. (E) Oxygen consumption rates (OCR) of the EMRA CD4^+^ and CD8^+^ T cells subsets were measured following a 15 minute stimulated with 0.5 μg/ml anti-CD3 and 5 ng/ml IL-2, the cells were then subjected to a metabolic stress test using the indicated mitochondrial inhibitors. Data are representative of 4 independent experiments. (F) The basal OCR, extracellular acidification rate (ECAR) and spare respiratory capacity were measured following a 15 minute stimulation with 0.5 μg/ml anti-CD3 and 5 ng/ml IL-2. Graphs show the mean ± SEM for 4 donors. (G) ATP concentration in EMRA T cell subsets, graphs show the mean ± SEM for 5 donors. p-values were calculated using a T-test.

We then examined differences in mitochondrial respiration between the CD4^+^ and CD8^+^ EMRA subsets. Differences were found in both the baseline respiration and respiration following injection of oligomycin, FCCP and rotenone and antimycin A (Figure 2E). The CD4^+^ EMRA population retain their ability to respond to challenge akin to the other CD4^+^ memory subsets (Supplementary Fig.2B), while we have shown this not to be the case for the CD8^+^ EMRA compartment^7^. The oxygen consumption rate (OCR), together with the spare respiratory capacity, the potential amount of stored energy a cell has to respond to challenge, were both upregulated in CD4^+^ EMRAs. Further implying a difference in mitochondrial content. While the extracellular acidification rate (ECAR), a marker of lactic acid production and glycolysis was only marginally increased compared to the CD8^+^ EMRA subset (Figure 2F). Furthermore the amount of ATP made by CD4^+^ EMRAs was also greater than that of the CD8^+^s (Figure 2G). These results suggest that the CD4^+^ EMRA subset have enhanced mitochondrial fitness that allows for a greater flexibility in the type of metabolism they can engage.

### CD8^+^ EMRA T cells display impaired nutrient uptake

T cells utilise a variety of energy sources including glucose and lipids, however their metabolic preferences are governed not only by their differentiation status but also by mitochondria fitness^16^. Indeed a lack of regulatory control over nutrient usage is a recurrent theme accompanying senescence and ageing^17^. We therefore sort to determine whether there were differences in glucose and fatty acid uptake in CD4^+^ and CD8^+^ EMRA T cell subsets. CD4^+^ EMRAs were found to take up more of the fluorescent glucose analogue 2-NBDG from their extracellular environment than their CD8^+^ EMRA counterparts (Figure 3A). Indeed CD4^+^ EMRAs showed higher expression of Glut-1, the major glucose transporter in T cells using an RNA-labelled probe (Figure 3B). Analysis of microarray data revealed high expression of alternate glut family members^18^. Interestingly CD8^+^ EMRAs displayed a higher expression of the class III glucose transporters Glut-8 and -10 (Figure 3B). Both these transporters are found intracellularly and are thought to transport glucose or galactose across intracellular organelle membranes^19^. The uptake of fluorescently labelled palmitate, BODIPY C16, a long chain fatty acid was also quantified. Similar to our observations for glucose uptake, CD4^+^ EMRA T cells also utilise significantly more palmitate than their CD8^+^ counterparts (Figure 3C). Furthermore CD4^+^ EMRA T cells express higher levels of both the fatty acid translocase CD36 and the fatty acid transporters FATP-2 and -3 (Figure 3D). Taken together these results suggest that the increased mitochondrial fitness of CD4^+^ EMRA T cells enable these cells to better utilise glucose and lipids, which may limit the impact of senescence.

**Figure 3.**
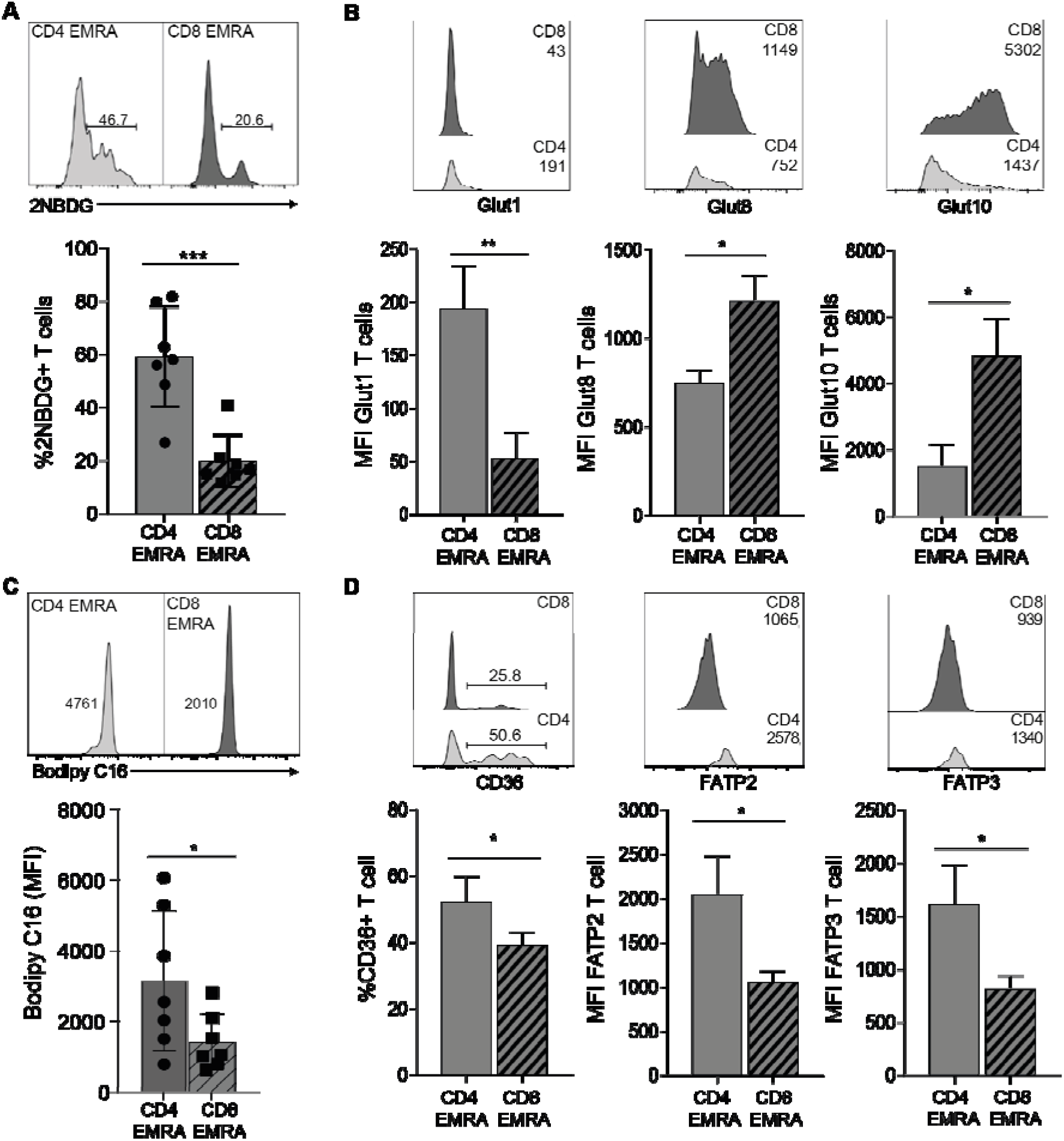
Impaired nutrient uptake by CD8^+^ EMRA T cells. (A) Glucose uptake was assessed using the fluorescent glucose analog 2-NBDG in CD4^+^ and CD8^+^ T CD45RA/CD27 defined EMRA T cells by flow cytometry following a 15 minute incubation. Data expressed as mean ± SEM of 7 donors. (B) Examples and data showing expression of the glucose transporters glut1 −8 and −10 in senescent T cells subsets directly *ex vivo*. Graphs show the mean ± SEM for 4 donors. (C) Lipid uptake was measured using fluorescently labelled palmitate, BODIPY C16 by flow cytometry following a 15 minute incubation in CD4^+^ and CD8^+^ EMRA T cells. Data expressed as mean ± SEM of 7 donors. (D) Examples and graphs showing the fatty acid translocase CD36 and FATP2 and −3 directly *ex vivo*. Data expressed as mean ± SEM of 6 donors. p-values were calculated using a T-test.

### Impaired proliferation and migration of CD8^+^ EMRA T cells

The end result of the DNA damage response is the activation of p53. p53 regulates cell cycle arrest limiting cell growth and proliferation, as well as playing a crucial role in limiting cell motility, a critical process for optimal T cell function^20^. In line with the theory that the acquisition of the CD4^+^ EMRA T cell subset occurs at a slower rate than their CD8^+^ counterpart, we find that the expression of p-p53 is higher in the CD8^+^ EMRAs compared to the CD4^+^s (Figure 4A). Although the expression of p-p53 in the CD4^+^ EMRAs is the highest of all the CD4^+^ memory subsets (data not shown). Furthermore the proliferative defect is more pronounced CD8^+^ EMRA subset, measured using ki67 (Figure 4B) and migration impaired (Figure 4C). Transwell chemotactic assays were used to assess migration, HUVECs were activated using 20% autologous donor sera, in order to create a more appropriate *ex vivo* environment and was found to be no different to activation with IFNγ (Supplementary Figure 3). Migration was assessed in response to CXCL10 and CXCL12, chemokines promoting the migration of memory T cells or 20% autologous serum. CD4^+^ EMRA T cells were less able to respond to CXCL10 and −12 than autologous serum, presumably due to the loss of CXCR3 and −4 from the CD4^+^ EMRA T cells^21, 22, 23^. CD8^+^ EMRAs on the other hand retain expression of CXCR3 and −4 and migrate in response to both chemokines and autologous serum, all be it to a lesser extent than their CD4^+^ counterparts (Figure 4C).

**Figure 4.**
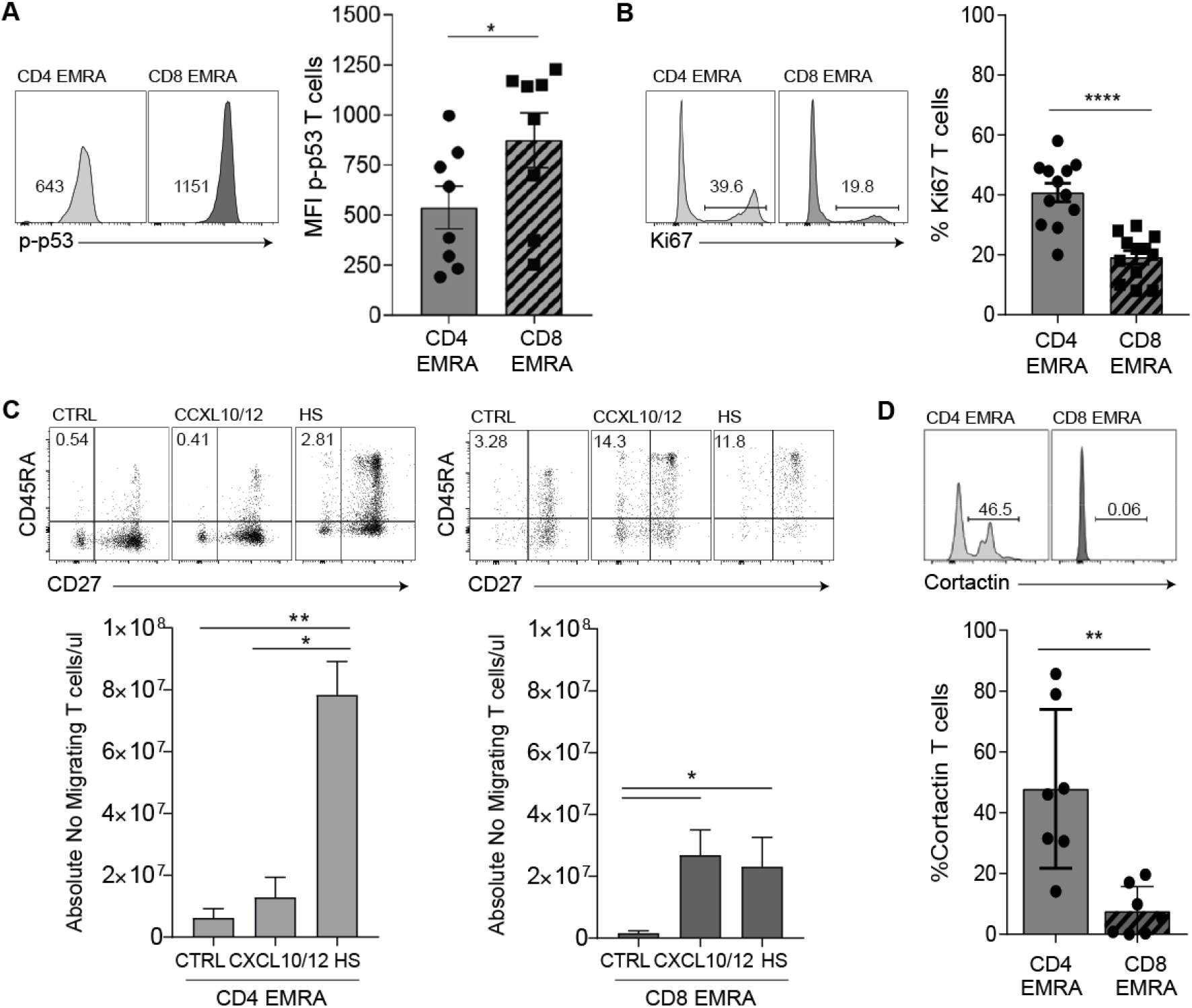
Impaired function observed in CD8^+^ EMRA T cells. (A) Example and graph showing the expression of p-p53 in CD4^+^ and CD8^+^ CD45RA/CD27 define EMRA T cells directly *ex vivo*. Graphs show the mean ± SEM for 4 donors. (B) Proliferation was defined in senescent T cell subsets using Ki67 directly *ex vivo*. Data show the mean ± SEM for 12 donors. (C) The migration of CD4^+^ and CD8^+^ EMRA T cells through HUVECs and their supporting transwell filters. HUVECs were stimulated with 20% decomplemented (heated at 56°C for 20min) autologous donor sera for 24h. PBMCs were allowed to adhere and migrate for 4h towards either media, CXCL10/12 or autologous serum. The number of T cells were counted and expressed as a percentage of the total CD4^+^ or CD8^+^ T cell subset added. Data are expressed as the mean ± SEM of 6 donors. (D) Representative flow cytometry plots and cumulative graphs showing the percentage of cortactin positive senescent T cell subsets analysed directly *ex vivo*. Data expressed as mean ± SEM of 7 donors. p-values were calculated using a T-test.

The enhanced migratory capacity of the CD4^+^ EMRA subset was also evident in their enhanced cortactin expression (Figure 4D). Cortactin is known to mediate complex roles in cell migration and invasion^24^; where it is involved in the formation of lamellipodia and invadiopodia^25, 26^. Furthermore the loss of p53 has been shown to promote invasion^27^. Taken together with the findings that p53 inhibits mitochondrial biogenesis, CD4^+^ EMRAs potentially retain more functionality than their CD8^+^ counterpart, as they are better able to undergo the necessary metabolic reprogramming needed to generate effector functions owing to their higher mitochondrial mass.

### Impairing mitochondrial function in CD4^+^ T cells accelerates senescence

We then wanted to investigate whether impairing mitochondrial function in CD4^+^ T cells could induce a similar phenotype observed in the CD8^+^ compartment. We used rotenone, a complex I inhibitor, to damage mitochondria. After a 5 day treatment with a low dose (10 nM) rotenone we observed an increased amount of low mass mitochondria at the expense of the higher more functional fused mitochondria in CD4^+^ T cells (Figure 5A). The change in mitochondrial state lead to a switch in metabolism, with the rotenone treated CD4^+^ T cells showing a strong response to the addition of glucose akin to the CD8^+^ T cells (Figure 5B). The rotenone treatment increased the basal glycolysis levels as well as increasing the glycolytic capacity of CD4^+^ T cells (Figure 5C). There is growing evidence that p53 can also regulate mitochondrial function, maintaining mitochondrial respiration through the transactivation of SCO2 (synthesis of cytochrome c oxidase 2)^28^. However stress conditions led to the translocation of p53 from the nucleus to the mitochondria leading to mitochondria mediated apoptosis^29^. We observe here that treatment of CD4^+^ T cells with rotenone leads to high levels of phospho-p53 (Figure 5D) and a slowing in cell growth that eventually lead to a loss of CD4^+^ T cells (Figure 5E). Therefore leading us to conclude that the higher mitochondrial mass observed in CD4^+^ T cells is protective against senescence by maintaining an oxidative state.

**Figure 5.**
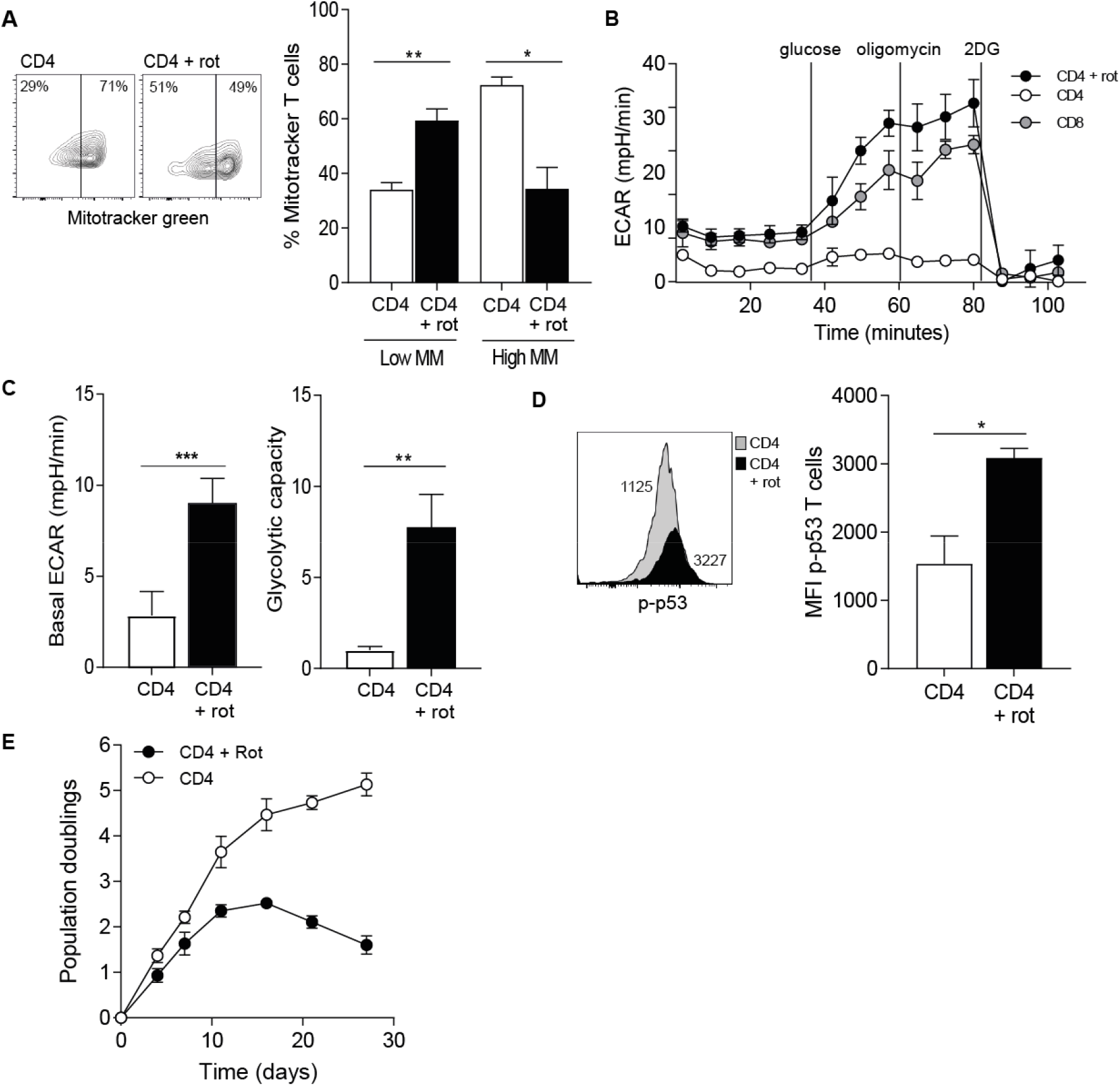
Impairing mitochondrial function in CD4^+^ T cells accelerates senescence. (A) Example and graph showing the mitochondrial mass of CD4^+^ T cells treated for 5 days with 10 nM rotenone or DMSO control. Graph show the mean ± SEM for 3 donors (B) Extracellular acidification rates (ECAR) of the rotenone or DMSO treated CD4^+^ T cells or DMSO treated CD8^+^ T cell were measured following a 15 minute stimulation with 0.5 μg/ml anti-CD3 and 5 ng/ml IL-2. The cells were then subjected to a glycolytic rate assay using the indicated substances. Data are representative of 3 independent experiments. (C) The basal ECAR and glycolytic capacity were measured following a 15 minute stimulation with 0.5 μg/ml anti-CD3 and 5 ng/ml IL-2. Graphs show the mean ± SEM for 3 donors. (D) Example and graph showing the expression of p-p53 in CD4^+^ T cells treated for 5 days with 10 nM rotenone or DMSO control. Graphs show the mean ± SEM for 3 donors. (E) Population doublings for CD4^+^ T cells treated 10 nM rotenone or DMSO control over 27 days. Graphs show the mean ± SEM for 3 donors.

## Discussion

Immunosenescence is a hallmark feature of ageing and is accompanied by a chronic low grade inflammatory state. Together these features are important drivers in numerous age-related pathologies^30^. Senescent or EMRA T cells are a highly dynamic and heterogeneous subset of cells that accumulate with age^18^ and are found in both the CD4^+^ and CD8^+^ T cell compartments. We and others have shown the CD8^+^ EMRA subset to accumulate more rapidly than their CD4^+^ counterparts with age^9, 31, 32^. Both subsets undergo the same phenotypic and functional changes, showing loss of co-stimulatory molecules and the acquisition of NK cell markers, together with a reduction in proliferative capacity and changes in cytokine production. It has been suggested that the CD4^+^ EMRA subset is more resistant to the effects of age owing to better homeostatic control in this compartment compared to that of CD8^+^ T cells. However we show here that it is enhanced mitochondrial dysfunction in the CD8^+^ EMRA subset that alters metabolic stability that governs the susceptibility of an immunosenescent state.

Mitochondrial dysfunction is a central event in many pathologies and contributes to age-related processes. Mitochondria have been shown to participate in every aspect of aging, such as a decline in stem cell function, cellular senescence and the development of the low grade inflammatory state^33^. Alterations that occur to mitochondria with age are numerous, including reductions in mitochondrial mass^34^, defects in mitochondrial biogenesis^35^, and impaired mitochondrial function in terms of ATP production and respiratory chain capacity^36^. Indeed we show here that the CD8^+^ EMRA T cell subset displayed a lower mitochondrial mass, impaired mitochondrial biogenesis, as well as having a hypopolarised mitochondrial phenotype compared to the CD4^+^ EMRA subset. When taken together these data indicate that the CD8^+^ EMRA subsets has a greater degree of mitochondrial impairment than the CD4^+^ compartment. We do not wish to oversimplify the idea that the presence of highly active mitochondria increases senescence resistance, as studies have demonstrated that mild reduction of mitochondrial function can counter-intuitively increase lifespan in lower organisms^37^. However the overexpression of mitochondrial enzymes in yeast increases lifespan and caloric restriction^38^, a well-established process to increase life span, mediates its effects through improved mitochondrial activity. Therefore we believe that the enhanced mitochondrial function observed for the CD4^+^ EMRA T cell subset does confer a survival advantage at the cellular level.

Increases in ROS levels have been demonstrated to be critical for the induction and maintenance of cell senescence^39^. However, ROS are also secondary messengers in cellular antioxidant pathways and rather than being thought of as deleterious by products and can be beneficial through the induction of an adaptive response that counteracts the rise in oxidative stress (mitohormesis)^40^. Indeed low levels of ROS have been implicated in improved cellular fitness and lifespan extension in various animal models^41^. The mitochondrial membrane potential (Δψm) is the central bioenergetic parameter controlling the generation of ROS^42^. We show here that the CD8^+^ EMRA subset displayed a hypopolarised phenotype whereas the CD4^+^ EMRAs were in a hyperpolarised state, with both subsets showed very little protein leak. As expected from cells that display a high Δψm CD4^+^ EMRAs also produced more ROS than their CD8^+^ counterparts. However the higher mitochondrial content observed in CD4^+^ EMRA T cells means these cells have more buffering capacity to quench the effects of the ROS. We postulate that the CD4^+^ EMRAs are better at controlling the damaging effect of ROS and are therefore better able to control the rate of senescence. Similar mechanisms have been reported in hepatocytes, whereby a reduction in mitochondrial ROS production, through a decrease in proton leak and a higher Δψm were found to be beneficial^43, 44^. However, it remains to be established what role uncoupling proteins (UCPs) play in mitigating ROS production by senescent T cells. Mouse models have shown a positive correlation between increased uncoupling and lifespan^45^. We can only infer a potential role for UCP1. UCP1 is activated by fatty acids and we show here that the CD4^+^ EMRA population take up more fatty acids and retain more fatty acid transporters than the CD8^+^ EMRA subset.

Mitochondria also play a key role in cellular metabolism, they house the electron transport chain and the TCA cycle, as well as playing crucial roles in the synthesis and breakdown of lipids^46^. Metabolic regulation plays an important role during cellular senescence, with dysregulated metabolism now identified as a feature of many different cell types including T cells^47^. There is increasing evidence that cell cultures become glycolytic as they age, depending on the senescence induction method used^47, 48^. Indeed we have shown previously that CD8^+^ EMRA T cells as they differentiate lose their metabolic plasticity and become more glycolytic^8^. However, we show here that CD4^+^ EMRAs retain their metabolic flexibility showing better glycolytic and oxidative capacity than their CD8^+^ counterparts. Furthermore, they also retain better glucose and lipid uptake together with increased expression of transporters. Interestingly CD8^+^ EMRAs displayed increased expression of the glucose transporters glut8 and −10. It is tempting to speculate in the light of a recent publication showing that glut6 functions as a glycolysis modulator in inflammatory macrophages without influencing glucose uptake^49^, that glut8 and −10 may act in a similar manner in senescent CD8^+^ T cells. Glut6, −8 and −10 are all members of the Glut III family of transporters, with glut6 and −8 being very closely located on chromosome 9 and glut10 having a very high affinity for both 2-deoxy-D-glucose and D-galactose^50^, both slow the rate of glycolysis. However further work needs to be carried out in order to establish whether glut8 and −10 play a negative role on glucose uptake on T cells.

The changes that occur to nutrient usage during senescence impacts T cell function, as is evident from the accelerated loss of function seen in CD8^+^ EMRA T cells. Senescent CD8^+^ T cells showed greater impairment in proliferative capacity than their CD4^+^ EMRA counterparts, in part, due to their reduced ability to take up and efficiently utilise metabolites. However the phenotype seen in CD8^+^ EMRA T cells also mirrors that reported for haematopoietic cells undergoing ER stress: compromised mitochondrial function, lower ATP levels and reduced glut1 expression^51^. Cell metabolism also exerts a strong influence on T cell migration^52^. The improved mobilisation of energy substrates by CD4^+^ EMRA T cells allows them to be better equipped to deal with the high energy demands of the migration process. Interestingly the CD4^+^ EMRAs were unable to respond to CXCL10 and −12, as the expression of CXCR3 and −4 have both been shown to decrease with differentiation in the CD4^+^ subset but is retained by CD8^+^s^21, 22, 23^. However, chemokine receptor expression in CD4^+^ T subsets does not universally decline, as the CD4^+^ EMRA subset show robust transmigration in response to autologous serum. The migratory advantage of CD4^+^ EMRA T cells may also be explained by their increased expression of cortactin, a core element of T cell locomotion involved in the formation of lamellipodia and invadiopodia^25, 26^. Taken together the greater loss in proliferative and migratory capacity of senescent CD8^+^ T suggests a potential greater impairment to CD8^+^ immunity. Indeed, failure to produce an antibody response following flu vaccination has been associated with an increase in senescent CD8^+^ T cells^53^.

We postulate here that mitochondrial density influences the extent of T cell senescence in a p53 dependent manner. Studies have shown that p53 can influence mitochondrial function, under steady state conditions p53 maintains mitochondrial respiration through the regulation of SCO2^28^. SCO2 is critical for regulating the cytochrome c oxidase (COX) complex, the major site of oxygen utilisation. However under stress settings p53 functions to control mitochondrial quality through the overexpression of SCO2, ROS generation and the removal of mitochondria^29^ via mitophagy or a protease-dependent degradation of damaged proteins^54^. We show here that the relatively lower expression of p53 in CD4^+^ versus CD8^+^ EMRA T cells maintains an oxidative metabolism, increasing mitochondrial stress in the CD4^+^ T cells leads to a more CD8^+^ like glycolytic metabolism that is accompanied by increased apoptosis. Our work goes against findings in fibroblasts where the targeted depletion of mitochondria through the impairment of their biogenesis demonstrated that decreased numbers of mitochondria were able to prevent the senescence response^55^. It is possible that for T cells, therapies aimed at increasing mitochondrial mass would be beneficial in combating the detrimental effects of senescence during ageing.

Collectively our results suggest that mitochondrial mass controls the senescence phenotype in T cells. However the mechanism remains elusive, it is not via a DNA damage response that has been suggested by others^55^. Mitochondria are a central part in inducing and governing the rate of T cell senescence. Therefore identifying T cell specific therapies aimed at increasing mitochondrial mass would be beneficial in combating the detrimental effects of senescence during ageing.

## Experimental Procedures

### Blood sample collection, isolation and cell culture

Heparinised peripheral blood samples were taken from healthy volunteers, average age 41 years ± 5. Healthy was taken as individuals who had not had an infection or immunisation within the last month, no known immunodeficiency or history of chemotherapy or radiotherapy, and were not receiving systemic steroids within the last month or any other immunosuppressive medications within the last 6 months. PBMC were isolated using Ficoll hypaque (Amersham Biosciences). All samples were obtained in accordance with the ethical committee of Royal Free and University College Medical School and the North East-York Research Ethics Committee 16/NE/0073. Human umbilical vein endothelial cells (HUVECs) were cultured according to the supplier’s instructions (Promocell)

### Flow cytometric analysis and cell sorting

Flow cytometric analysis was performed using the following antibodies: CD4 PECF594 (RM4-5) from BD Biosciences and CD8 PerCP (SK1), CD45RA BV605 (HI100), CD27 BV421 (O323), CD28 BV785 (CD28.2), CCR7 PECy7 (G043H7), CD36 APCCy7 (5-271) from BioLegend. FATP2 (Abcam) and FATP3 (Atlas antibodies) were measured in conjunction with goat anti-rabbit AF488 (Abcam). Cortactin expression was assessed using rabbit anti-human cortactin antibody (PA5-27134; Life Technologies) stained in conjugation with goat anti-rabbit Cy3 (Life Technologies). PGC1α (3G6), p-p53 (16G8) both from Cell Signaling, and Ki67 (B56; BD Bioscience) were assessed by intracellular staining using solution AB (ThermoFisher) and goat anti-rabbit AF488 (Abcam). All samples were run using an LSR II (BD Biosciences) and analysed using FlowJo software (Treestar).

Magnetic beads were used to isolation of CD8^+^ and CD4^+^ T cells (Miltenyi Biotec) according to the manufacturer’s instructions. The purity of T cell subsets was assessed by flow cytometry.

### PrimeFlow RNA assay

PrimeFlow RNA Assay technology was used to determine SLC2A gene expression according to the manufactures instructions (ThermoFisher). PBMCs were incubated with the following gene-specific probes: SLC2A10 (AF647 - Type 1 probe set), SLC2A8 (AF488 - Type 4 probe set), SLC2A1 (AF750 - Type 6 probe set). Samples were analysed immediately as described above.

### Proliferation assays

CD45RA/CD27 sorted CD4^+^ and CD8^+^ T cells were stimulated with 0.5μg/ml plate coated anti-CD3 (OKT3) and 5 ng/ml IL-2 for 3 days and proliferation was assessed by staining for the cell cycle related nuclear antigen Ki67 as described above.

### Transmission electron microscopy studies

CD27/CD45RA defined CD4^+^ and CD8^+^ EMRA T cell subsets were isolated and fixed in 2% paraformaldehyde, 1.5% glutaraldehyde in 0.1 m phosphate buffer pH 7.3. They were then osmicated in 1% OsO4 in 0.1M phosphate buffer, dehydrated in a graded ethanol-water series, cleared in propylene oxide and infiltrated with Araldite resin. Ultra-thin sections were cut using a diamond knife, collected on 300 mesh grids, and stained with uranyl acetate and lead citrate. The cells were viewed in a Jeol 1010 transmission electron microscope (Jeol) and imaged using a Gatan Orius CCD camera (Gatan). Mitochondrial volume density (percentage of T cell volume occupied by mitochondria) was determined from EM images using a point-counting method using image J.

### Mitochondrial measurements

Mitochondrial mass was assessed by incubating labelled PBMCs with 100 nM of MitoTracker Green FM (ThermoFisher) for 30 minutes at 37°C, 5% CO_2_. Mitochondrial membrane potential was investigated using TMRE (ThermoFisher), 1 μM TMRE was incubated with labelled PBMCs for 30 minutes at 37°C, 5% CO_2_. Mitochondrial ROS was measured using MitoSOX (ThermoFisher), 2μM MitoSOX was incubated with labelled PBMCs for 20 minutes at 37°C, 5% CO_2_. Unfixed samples were immediately collected on a LSR II (BD Bioscience).

### ATP determination

Intracellular ATP levels were measured ex vivo on sorted CD4^+^ and CD8^+^ T cell subsets via a bioluminescence assay according to the manufactures instructions (ThermoFisher).

### Metabolic assays

Oxygen consumption rates (OCR) and extracellular acidification rates (ECAR) were measured in CD45RA/CD27 sorted CD4^+^ and CD8^+^ T cell subsets following 15 minute stimulation with 1 μg/ml anti-CD3 and 5 ng/ml IL-2. The assay was performed in RPMI without phenol red and carbonate buffer (Sigma) containing 25 mM glucose, 2 nM L-glutamine and 1 mM pyruvate. The metabolic stress test was performed using 1 μM oligomycin, 1.5 μM fluorocarbonyl cyanide phenylhydrazone (FCCP), 100 nM rotenone and 1 μM antimycin A (Sigma) with the XF-96 Extracellular Flux Analyzer (Agilent). Glycolysis was examined via extracellular acidification rate (ECAR) in assay buffer excluding glucose, following injection with 10 mM glucose, 1 μM oligomycin and 100 mM 2-deoxy-D-glucose (2-DG; Sigma).

### Glucose and lipid uptake assays

To assess glucose and lipid uptake in T cell subsets, PBMCs were incubated with anti-CD3 (1 μg/ml) for 30 min at 37°C. Cells were subsequently incubated with 1nM Bodipy FL C_16_ or 100μM 2-NBDG, both from ThermoFisher in PBS and incubated for 15 minutes in media containing no glucose or serum. Samples were then analysed by flow cytometry.

### Transwell migration assay

HUVECs monolayers were grown to confluence on transwell membranes (Corning) in the presence of 20% autologous donor sera or 10 ng/ml IFNγ (R&D Systems). PBMCs from healthy donors were placed in M199 medium (Sigma) in the top well and the chemoattractant in the bottom well, which was either 20% autologous donor sera or 1 g/ml CXCL10 and CXCL12 (R&D Systems). Cell migration was assessed after incubation at 37°C for 4h, each condition was set up in duplicate transwells. Migrated T cells were then collected from the top and bottom wells respectively, stained with phenotypic markers and quantified for a fixed period of time (3 min) by flow cytometer. Counting beads (BD Bioscience) were also run to enumerate the total number of cell to have transmigrated.

### Rotenone cultures

Purified CD4^+^ T cells were incubated for 5 days with 10 nM rotenone (Sigma) or DMSO control after which time the cells were used for metabolic assessment: mitochondrial mass, p53 levels and extracellular flux analysis as describe above. For long term rotenone cultures, 10 nM rotenone was plated together with 0.2 × 10^6^ CD4^+^ T cells and 0.5 μg/ml anti-CD3 and 5 ng/ml IL-2. Cells were counted every 4/5 days and population doubling was calculated using the following equation: PD = log10(Nf/Ni)/log(2), Nf = number of cells harvested, Ni = initial cell number seeded.

### Statistical analysis

GraphPad Prism was used to perform statistical analysis. Statistical significance was evaluated using the paired Student *t*-test. Differences were considered significant when *P* was < 0.05.

