## Supplementary figures for "Mitochondrial mass governs the extent of T cell senescence"

**Supplementary figure legends**

**Supplementary Figure 1. Experimental gating strategy**

All experiments were first gated on the lymphocyte population, followed by live cells then the CD4^+^ or CD8^+^ population before the CD45RA/CD27 profile was obtained.

**Supplementary Figure 2. Mitochondrial changes to CD4^+^ T cell subsets.**

A) Mitotracker green staining in CD4^+^ CD27/CD45RA defined T cells analysed directly *ex vivo*. Data expressed as mean ± SEM of 6 donors. B) Oxygen consumption rates (OCR) of CD27/CD45RA defined CD4^+^ T cells were measured following a 15 minute stimulated with 0.5 µg/ml anti-CD3 and 5 ng/ml IL-2, the cells were then subjected to a metabolic stress test using the indicated mitochondrial inhibitors. Data are representative of 4 independent experiments.

**Supplementary Figure 3. Impaired migration of CD8^+^ T cells through HUVEC stimulated with chemokine**

The migration of CD4^+^ and CD8^+^ EMRA T cells was also tested through HUVECs that had been stimulated with 10 ng/ml IFNγ rather than autologous donor sera. PBMCs were allowed to adhere and migrate for 4h towards autologous serum. The number of T cells were counted and expressed as a percentage of the total CD4^+^ or CD8^+^ T cell subset added. Data are expressed as the mean ± SEM of 4 donors.

**Supplementary Figure 1.**

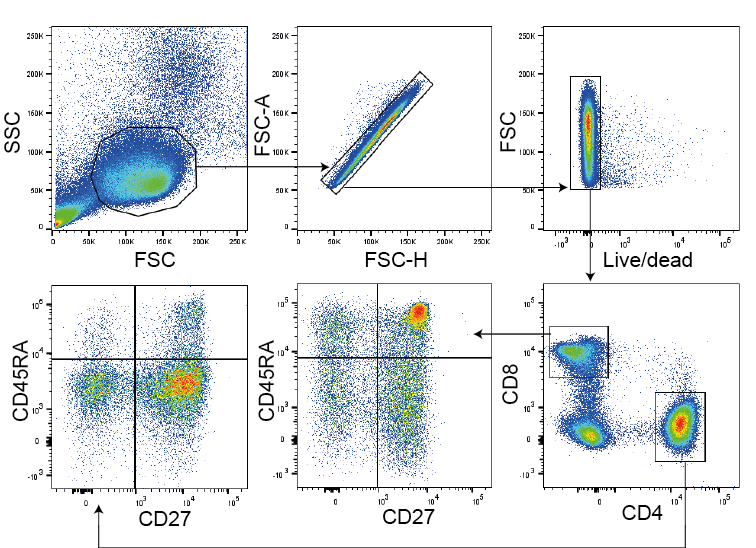

**Supplementary Figure 2.**

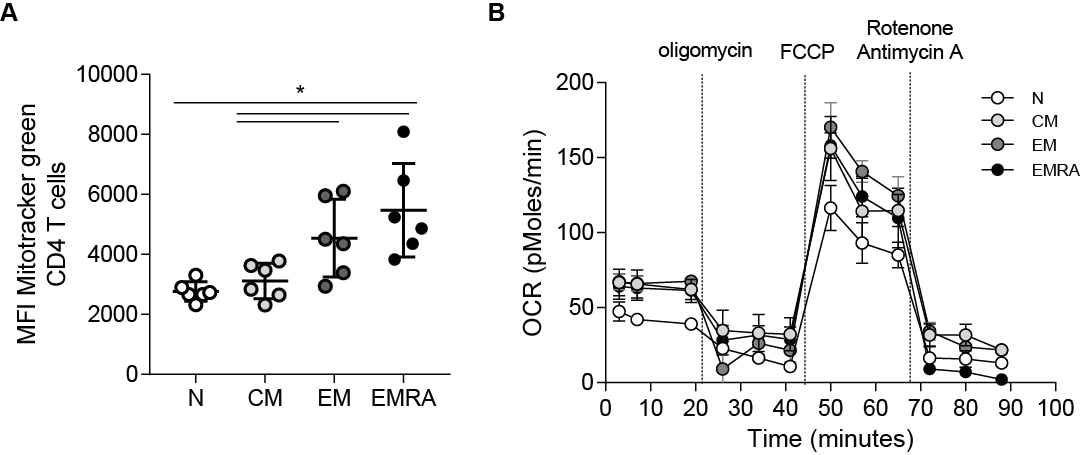

**Supplementary Figure 3.**

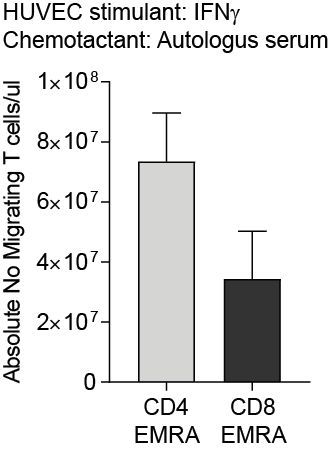
